# Functional High Throughput Drug Screening Reveals Cyproheptadine as a Novel Treatment for *LMNA*-related Cardiomyopathy

**DOI:** 10.64898/2026.06.12.731852

**Authors:** Hananeh Fonoudi, Ali Negahi Shirazi, Hui-Hsuan Kuo, Carlos G. Vanoye, Xiaozhi Gao, Susan Doody, Mariam Jouni, Brian Lenny, Achal Neupane, Janavi Kotamarthi, Yadav Sapkota, Jane E. Wilcox, Alfred L. George, Paul W. Burridge

## Abstract

**Objectives:** To define shared and variant-specific mechanisms underlying LMNA-associated dilated cardiomyopathy (DCM) and identify therapeutic candidates using human stem cell–based models.

**Background:** Variants in the gene LMNA, encoding lamin A/C, cause 5–10% of dilated cardiomyopathies (DCM) and are strongly associated with heart failure and arrhythmias. Yet, the mechanisms by which LMNA variants drive disease and the distinction between shared and variant-specific phenotypes remain unclear.

**Methods:** To address this, we generated human induced pluripotent stem cell-derived cardiomyocytes (hiPSC-CMs) from six LMNA-DCM patients carrying three pathogenic variants (T150A, E381Afs*39, R527H) and from five healthy control patients.

**Results:** LMNA hiPSC-CMs exhibited nuclear membrane deformation, reduced beat rate, arrhythmias, and prolonged calcium transients. Transcriptomic and electrophysiological analyses revealed downregulation of cardiac genes and ion channels, with abnormal Ca²⁺ handling emerging as a shared disease mechanism. Leveraging a high-throughput functional assay, we performed an unbiased drug screen and identified cyproheptadine, an FDA-approved antihistamine, as the only compound to alleviate abnormal function across all LMNA variants.

**Conclusion:** Our findings reveal a shared disease mechanism across multiple LMNA variants driven by dysregulated Ca²⁺ handling. This work establishes a patient-specific drug discovery platform and identifies cyproheptadine as a promising therapeutic candidate for LMNA-associated dilated cardiomyopathy.

**Highlights:**

- Patient-specific LMNA hiPSC-cardiomyocytes robustly recapitulate disease phenotypes, including nuclear defects, arrhythmias, and contractile dysfunction.
- Dysregulated calcium handling emerges as a unifying mechanism driving pathology across distinct LMNA variants.
- Variant-resolved analysis reveals both shared and mutation-specific molecular and functional signatures.
- High-throughput screening identifies cyproheptadine as a potent, broadly effective rescue agent across all tested LMNA variants.
- This work establishes a scalable precision medicine platform for rapid therapeutic discovery in inherited cardiomyopathies.

## Introduction

Lamin A and C (collectively referred to as lamin A/C) represent the primary protein isoforms encoded by the gene *LMNA*. Mutations in *LMNA* give rise to a broad spectrum of diseases collectively referred to as laminopathies, a group of disorders characterized by heterogeneous effects on the various tissues, including striated muscles, peripheral nerves, and adipose tissue. Among the diverse array of symptoms associated with these disorders, *LMNA*-related dilated cardiomyopathy (LMNA-DCM) is the most prevalent phenotype, leading to arrhythmias, ventricular dilation, and eventual heart failure. Variants in *LMNA* account for 5-10% of DCM cases, making it one of the most common known causes of this condition while also being associated with a comparatively poor prognosis. In families with DCM and conduction system abnormalities, this percentage of cases with *LMNA* variants is substantially higher (30-45%)^1,2^. While the age of onset and presentation of DCM symptoms may vary among patients, LMNA-DCM becomes symptomatic universally after age 60 years old^3–5^. The molecular pathways linking *LMNA* variants and cardiomyopathy remain incompletely understood, posing challenges in correlating specific *LMNA* variants with disease phenotypes and patient outcomes. Additionally, it is unclear which disease-causing pathways are common among LMNA-DCM patients, while others are variant-specific. The clinical phenotype is often progressive compared to other genetic causes of DCM and can be unpredictable with high risk of sudden cardiac death due to fatal arrhythmia. Unfortunately, no specific therapy currently exists for LMNA-DCM. Moreover, limited knowledge is available regarding the correlation between variants within specific *LMNA* domains and clinical outcomes, resulting in treatment strategies that are predominantly symptom oriented.

The structural organization of lamin A/C encompasses a head domain, followed by coil segments 1A, 1B, 2, and a tail domain. The varying penetrance of disease-related *LMNA* variants suggests the likelihood of domain-specific effects. Lamin A/C serves three major roles within the cell: firstly, it mediates the structural linkage between the nucleus and the cytoskeleton^6^; secondly, it interacts with chromatin and transcription factors to modulate gene expression^7^; and thirdly, it provides a platform for the assembly of complexes involved in signal transduction^8^. Consequently, any alteration in lamin A/C function has the potential to influence the resistance of the nuclear membrane to mechanical stresses^9,10^. This is of particular significance in mechanically strained tissues such as the heart. The relevance of these findings is underscored by observations in *LMNA* knockout mouse models, which recapitulate several features of human LMNA-DCM. These models exhibit elevated nuclear deformation and sensitivity to mechanical stress, and a propensity for arrhythmias^9,11,12^. Furthermore, *LMNA* knockout mice exhibit upregulation of the ERK, mTOR, and PDGF signaling pathways^13–15^. The same pathways have also been linked to *LMNA* variants^16^, suggesting a potential role for these pathways in the pathogenesis of LMNA-DCM. Remarkably, this phenotypic presentation can be reversed through the overexpression of *LMNA* in cardiomyocytes, underscoring a cell-autonomous component^17^. Based on these observations, therapeutic agents targeting the ERK, mTOR, and PDGF pathways are currently under investigation. Despite the advantages of using animal models, the phenotype in animal models can differ substantially from the phenotype in humans^18^*. LMNA* knockout mouse models exhibit severe and often exaggerated phenotypes that do not fully recapitulate the spectrum of human disease. In contrast, only a small number of mouse models harboring *specific LMNA* point mutations such as N195K^19^, H222P^20^, M371K^21^, and L530P^22^. have been generated. Together, these limitations underscore the need for human-based disease models to study LMNA-associated cardiomyopathy.

Human induced pluripotent stem cell-derived cardiomyocytes (hiPSC-CMs) been consistently shown to recapitulate key aspects of LMNA-DCM *in vitro* (reviewed in^23^), particularly blebbing and nuclear damage^24–26^. However, these investigations often encountered limitations due to the small number of hiPSC lines studied and a predominant focus on assessing the effects of individual *LMNA* variants. To comprehensively elucidate both common and variant-specific phenotypes and to establish correlations between *in vitro* phenotypes and underlying altered pathways, we harnessed patient-specific hiPSCs derived from individuals with LMNA-DCM, each harboring distinct *LMNA* variants residing in different domains. Our analysis confirmed existing findings and unveiled a common arrhythmogenic electrophysiological feature in all laminopathy hiPSC-CMs. Based on these observations, we developed an *in vitro* functional drug-response assay to execute a large-scale, unbiased, high-throughput drug repurposing screen. The primary goal was to identify novel compounds capable of mitigating the LMNA-DCM phenotype. Our functional analysis revealed specific drugs which were able to rectify the abnormal phenotype present in each of the studied laminopathy hiPSC-CMs. Notably, one compound, cyproheptadine, emerged as a shared therapeutic candidate across all the studied hiPSC-CMs. This places cyproheptadine as a strong candidate for further exploration in clinical trials, offering the promise of a potentially novel treatment approach for LMNA-DCM.

## Methods

The data, analytical methods, and study materials are available from other researchers upon reasonable request for purposes of reproducing the results. Detailed methods are provided in the Supplementary Information online. RNA–seq data have been deposited in Gene Expression Omnibus (https://www.ncbi.nlm.nih.gov/geo/) with accession code GSE320580.

### Patient specific human induced pluripotent stem cell culture and cardiac differentiation

Six volunteers diagnosed with dilated cardiomyopathy and a pathogenic variant in *LMNA* were enrolled in the study over the course of two years along with five healthy individuals were enrolled as controls. Protocols were approved by the Northwestern University Institutional Review Board (IRB STU00205725) and written informed consent was obtained from all volunteers. This study was conducted in accordance with the principles outlined in the Declaration of Helsinki. Blood samples were taken from each volunteer and peripheral blood mononuclear cells were isolated and reprogrammed to hiPSCs using a CytoTune-iPS 2.0 (Invitrogen, A16518) in the low-cost variant of B8 as previously described^27,28^. One hiPSC line from each individual was nucleofected with a plasmid containing exogenous *TNNT2* promoter-driven neomycin resistance cassette targeted to the AAVS1 locus (Addgene, 214013) to allow cardiomyocyte purification during differentiation and was used for downstream experiments. Whole genome sequencing was conducted on hiPSCs to verify the presence of LMNA variants and the absence of other pathogenic variants introduced during reprogramming. hiPSCs were differentiated into cardiomyocytes using the previously described RBAI protocol^29^. hiPSC-CMs were then purified using neomycin exposure and treated with complete maturation medium (MMc) to enhance their maturation level^30^.

### Calcium transient measurement

Intracellular calcium transients were recorded using an IC200 Kinetic Image Cytometer (KIC, Vala Sciences) and Cal-520 AM calcium dye and analyzed using CyteSeer software as previously described^31^. Briefly, purified hiPSC-CMs were dissociated on differentiation day 22 and plated at 100,000 cells per well in a Matrigel-coated 96-well black µClear microplates (Greiner, 655090). On differentiation day 30, cells were treated with Hanks’ Balanced Salt Solution (HBSS, Corning, 21-023-CV) supplemented with 20 mM HEPES (Hyclone, SH30237.01), 0.04% Pluronic F-127 (Sigma, P2443), 2.5 mM probenecid (Sigma, P8761), 2 drops per 10 mL NucBlue Live ReadyProbes Reagent (Invitrogen, R37605), and 2 μM Cal-520 AM (AAT Bioquest, 21130) for 1 h at 37 °C, 5% CO_2_. For imaging, Ca^2+^ dye loading solution cells were transferred to FluoroBrite DMEM (Gibco, A1896702) medium containing 2.5 mM probenecid. Calcium transients were recorded at 37 °C, 5% CO_2_ for a 10 s time series at an acquisition frequency of 99.75 Hz. The average of all calcium transients for each cell in the well was analyzed using CyteSeer Analysis software (Vala Sciences).

### Impedance analysis

Impedance recording was measured using a CardioExcyte96 multielectrode array (Nanion Technologies). Briefly, purified hiPSC-CMs were dissociated on differentiation day 22 and plated at 100,000 cells per well in a Matrigel-coated 96-well Cardioexcyte96 sensor plates with stimulation electrode (Nanion, 20-1003). At differentiation day 30 impedance of spontaneously beating cells was recorded every 5 min for at least 1 h using a 30 sec sweep duration. Data was analyzed using Nanion Data Control software. During recording cells were maintained at 37 °C, 5% CO_2_, and 80% humidity.

### CRISPR/Cas9-Mediated Homology-Directed Repair

Correction of the three *LMNA* variants analyzed in this study was performed in LMNA3, LMNA5, and LMNA7 hiPSC lines. Twenty–base pair guide RNAs and custom guide-specific Alt-R CRISPR-Cas9 crRNAs were designed using the IDT design tool. Alt-R CRISPR-Cas9 crRNA and Alt-R CRISPR-Cas9 tracrRNA (Integrated DNA Technologies, IDT) were reconstituted in nuclease-free duplex buffer at 200 µM. crRNA and tracrRNA were combined at a 1:1 molar ratio (final duplex concentration: 100 µM), heated to 95 °C for 5 min, and cooled to room temperature. The duplex was then complexed with Alt-R S.p. HiFi Cas9 Nuclease V3 (IDT, 1081061) at a 1:2 molar ratio (Cas9:RNA duplex) and incubated for 20 min at room temperature to allow ribonucleoprotein (RNP) complex formation. hiPSCs were dissociated with TrypLE (3 min, room temperature), and 1 × 10^6^ cells were electroporated with the RNP complex and Alt-R Electroporation Enhancer (IDT, 1075916; final concentration: 100 µM). To evaluate crRNA cutting efficiency, pooled cells were harvested four days post-transfection, and genomic DNA was extracted for Sanger sequencing. Editing efficiency was quantified using TIDE analysis (https://tide.nki.nl/). The single- stranded oligodeoxynucleotide (ssODN) repair template used for CRISPR/Cas9-mediated homology-directed repair (HDR) was 90 nucleotides in length and included a silent mutation to facilitate clone identification. hiPSCs were co-transfected with Cas9, the RNP complex, the repair template (4 µM), electroporation enhancer, and HDR Enhancer V2 (1 µM; IDT, 10007921) containing validated crRNA. Transfected cells were plated at low density (distributed across six wells of a 6-well plate). Individual clones were manually isolated 3–5 days post-transfection using a P20 pipette. At the subsequent passage, clones were screened by Sanger sequencing to confirm successful editing. Primer sequences used for Sanger validation are provided in **Supplemental Table 1**.

### Drug screening

Purified hiPSC-CMs were dissociated at differentiation day 22 and plated at 25,000 cells per well in a Matrigel-coated 384-well black µClear microplates (Greiner, 781091). On differentiation day 27, the medium was replaced with RPMI 1640 without L-glutamine and phenol red (Corning, 17-105-CV) supplemented with 500 mg/ml recombinant human serum albumin (Oryzogen, HYC002M01) and GlutaMAX supplement (Thermofisher, 35050061). Cells were then exposed to the Prestwick Chemical Library®—1280 compounds using a LabCyte Echo 550 liquid handling instrument (Beckman Coulter) at a final concentration of 1 µM. After 72 h cells were assessed for their calcium transients using an IC200 KIC.

### Statistical Analysis

Data was analyzed in Excel and GraphPad Prism 10. Detailed statistical information is included in the corresponding figure legends. All data were presented as mean ± SEM. Data was checked for normal distribution and comparisons were conducted using ordinary one-way ANOVA and normalized based on multiple tests. Significant differences were defined as *P* < 0.05 (*), *P* < 0.01 (**), *P* < 0.001 (***), and *P* < 0.0001 (****).

## Results

### Generation of patient-specific hiPSCs from laminopathy families

Six volunteers with pathogenic variants in *LMNA* and a history of LMNA-DCM were recruited from four families (**Fig. 1A**). Among the six patients, two individuals (LMNA1 and LMNA2 from family 1) had pathogenic variants in coil1B (c.488A>G, p.T150A). Two individuals from family 2 (LMNA3 and LMNA4) presented a frameshifting deletion in coil2 (c.1142-1157+1del17, p.E381Afs*39). Two remaining individuals (LMNA5 from family 3 and LMNA6 from family 4) harbored a pathogenic *LMNA* variant in the tail domain (c.1580G>A, p.R527H) (**Fig. 1B**). All detected variants were heterozygous. Five unrelated healthy volunteers with no history of cardiac disease were used as controls. hiPSC lines were established from peripheral blood mononuclear cells (PBMCs) using non–integrating reprogramming and our well–established protocols^27–29,32^. These lines showed normal hiPSC morphology and expressed high levels of undifferentiated markers POU5F1, SOX2, SSEA4, and TRA-1-60 (**Supplementary Fig. 1A** and **B**). Whole genome sequencing was used to verify the presence of expected *LMNA* variants (data not shown). hiPSCs were differentiated into cardiomyocytes using our established chemically defined monolayer protocol^29^ which consistently produced >90% pure cardiomyocytes. hiPSC-CMs were later treated with a maturation media to enhance the maturation of cardiomyocytes^30^.

**Figure 1.**
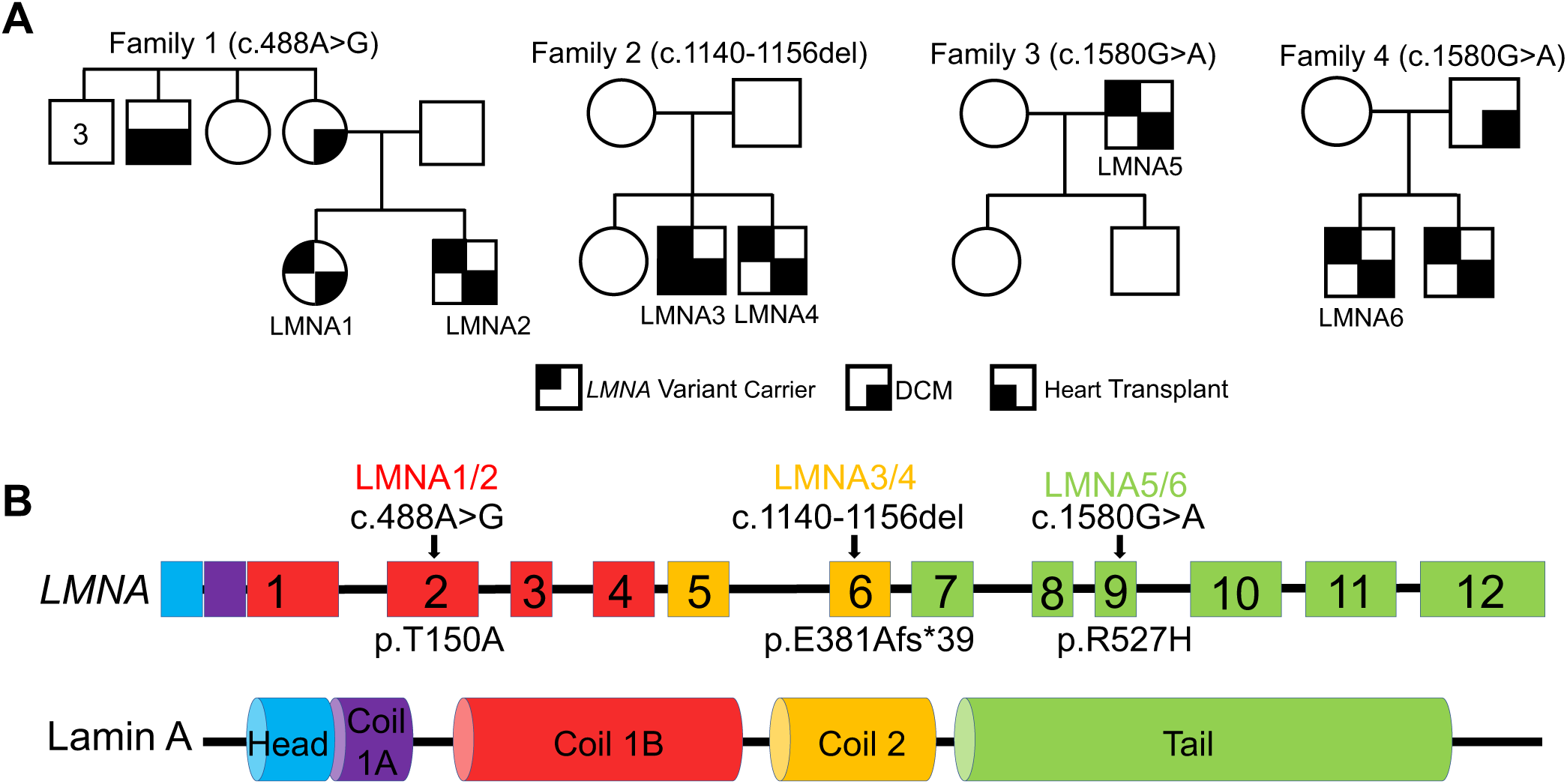
Family pedigrees and schematic diagram of *LMNA* gene and protein. **A)** Family pedigrees of all the individuals in the study. Circle and square indicate female and male individuals respectively. **B)** Summary of the *LMNA* variants and their corresponding gene and protein location. The code for each individual (LMNA1-6) is shown in the diagram based on the location and codon changes of the variant present in each individual. Below the gene panel the corresponding protein change for each variant is shown.

### Patient-specific hiPSC-CMs recapitulate nuclear deformation observed in LMNA-DCM

We first assessed the morphology of hiPSC-CMs nuclear membranes as it has previously been demonstrated that hiPSC-CMs with a *LMNA* variants (R225X but not Q354X and T518fs) have an abnormal nuclear morphology called nuclear blebbing, attributed to the role of Lamin A/C in maintaining nuclear membrane integrity, which increases in frequency after electrical stimulation^24^. Lamin A/C immunofluorescent staining of hiPSC-CMs (**Fig. 2A**) showed a higher frequency of the blebbing phenotype in all six of our *LMNA* variant hiPSC-CMs when compared to control hiPSC-CMs (**Fig. 2B**). These findings underscore the pivotal role of Lamin A/C in maintaining nuclear membrane integrity and highlight the validity of hiPSC-CMs as an *in vitro* model system to faithfully recapitulate the laminopathy cellular phenotype.

**Figure 2.**
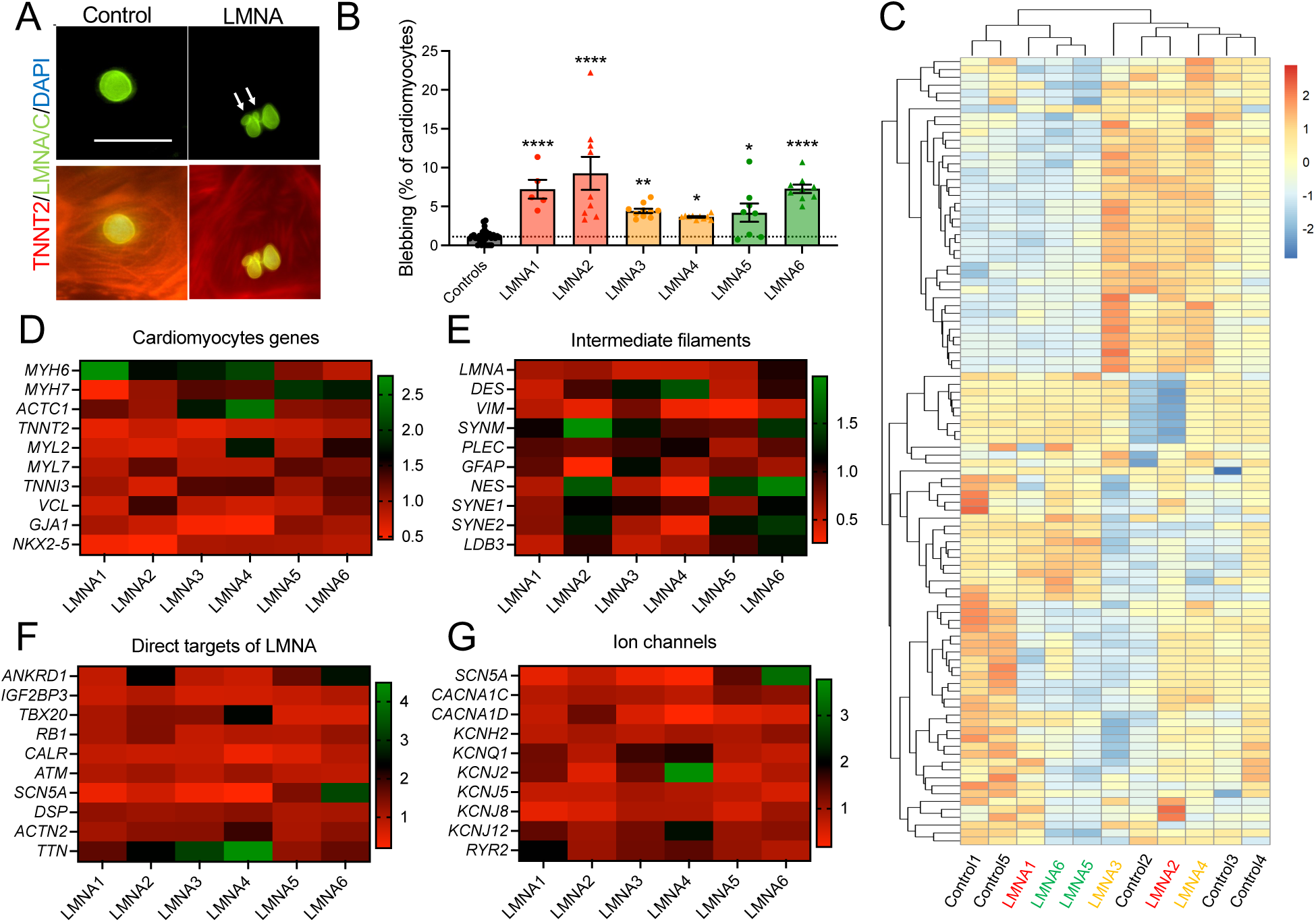
Assessment of nuclear membrane and transcriptome analysis of *LMNA* hiPSC-CMs. **A)** Representative images of control and laminopathy hiPSC-CMs stained with TNNT2 and LMNA/C. Scale bar is 200 µm. **B)** Quantification of the nuclear membrane deformation in controls and *LMNA* hiPSC-CMs using ImageJ software. Each dot represents the number of assessed fields of view. Control N = 30, LMNA1 N = 5, LMNA2 N = 9, LMNA3 N = 10, LMNA4 N = 10, LMNA5 N = 8, LMNA6 N = 9. The dotted line indicates the value for the average of five control hiPSC-CMs. Data is analysed using one way ANOVA. **** indicate *P-*value < 0.0001. **C)** Unsupervised hierarchical clustering of control and laminopathy hiPSC-CMs. Heat maps indicating the gene expression of **D)** cardiac markers, **E)** intermediate filaments, **F)** direct targets of LMNA and **G)** ion channels, from each laminopathy patient compared to controls. All gene expression data is normalized to the average of controls.

### Gene expression signature in LMNA-DCM hiPSC-CM is altered

To harness the gene expression profile change in LMNA-DCM, the transcriptome of six laminopathy patients and five control hiPSC-CMs was assessed (**Fig. 2C**). Pathway enrichment analysis of the differentially expressed genes revealed presence of cardiomyopathy related pathways (**Supplementary Fig. 2**). This finding was further supported by the lower expression of the genes involved in the cardiomyocytes structure and function including *MYH6, MYH7, ACTC1, TNNT2, MYL2, MYL7, TNNI3, VCL, GJA1*, and *NKX2-5* compared to controls (**Fig. 2D**). Interestingly, expression of *LMNA* itself was significantly lower among all the laminopathy hiPSC-CMs while other intermediate filament genes including *DES, VIM, SYNM, PLEG, GFAP, NES, SYNE1, SYNE2,* and *LDB3* did not show similar common pattern (**Fig. 2E**), suggesting a role for the variants within *LMNA* in controlling its own gene expression. To further assess the effect of downregulation of *LMNA*, the expression of genes encoding proteins that are direct targets of LMNA (*ANKRD1, IGF2BP3, TBX20, RB1, CALR, ATM, SCN5A, DSP, ACTN2*, and *TTN*) (**Fig. 2F**) were studied. An overall downregulation of these genes was observed indicating a potential role for *LMNA* in controlling their expression and subsequently the observed abnormal phenotype in laminopathy hiPSC-CMs. Finally, pathway enrichment analysis revealed abnormal Ca^2+^ handling in *LMNA* hiPSC-CMs, suggesting a role for Ca^2+^ related pathways in arrythmia associated with LMNA-DCM (**Supplementary Fig. 2B**). Similarly, when expression of ion channel genes (*SCN5A, CACNA1C, CACNA1D, KCNH2, KCNQ1, KCNJ2, 5, 8, 12,* and *RYR2*) was assessed, *LMNA* hiPSC-CMs exhibited lower expression compared to controls (**Fig. 2G**).

### Electrophysiological properties of laminopathy hiPSC-CMs are abnormal

To further assess the arrhythmogenic features of laminopathy hiPSC-CMs, we performed patch clamp recordings to assess electrophysiological properties. Manual current clamp recordings performed on two *LMNA* variant carrier lines (LMNA3 and 4) revealed a common signature of spontaneous arrhythmia and early after depolarization (**Fig. 3A**). Additionally, the duration of evoked action potentials (APD) was prolonged in both LMNA3 and 4 compared to control (**Fig. 3B)**. More specifically, *LMNA* hiPSC-CMs exhibited significantly slower maximal upstroke velocity (V_max_) (**Fig. 3C**). APD at 30% and 50% completion were not significantly different between *LMNA* hiPSC-CMs and controls while a significant prolongation was detected at 90% completion of (**Fig. 3D**). These data could be consistent with an increased/non-inactivating inward current, or a decrease in an outward current^33^, which suggests the involvement of Na^+^ and Ca^2+^ channels in arrythmia associated with LMNA-DCM.

**Figure 3.**
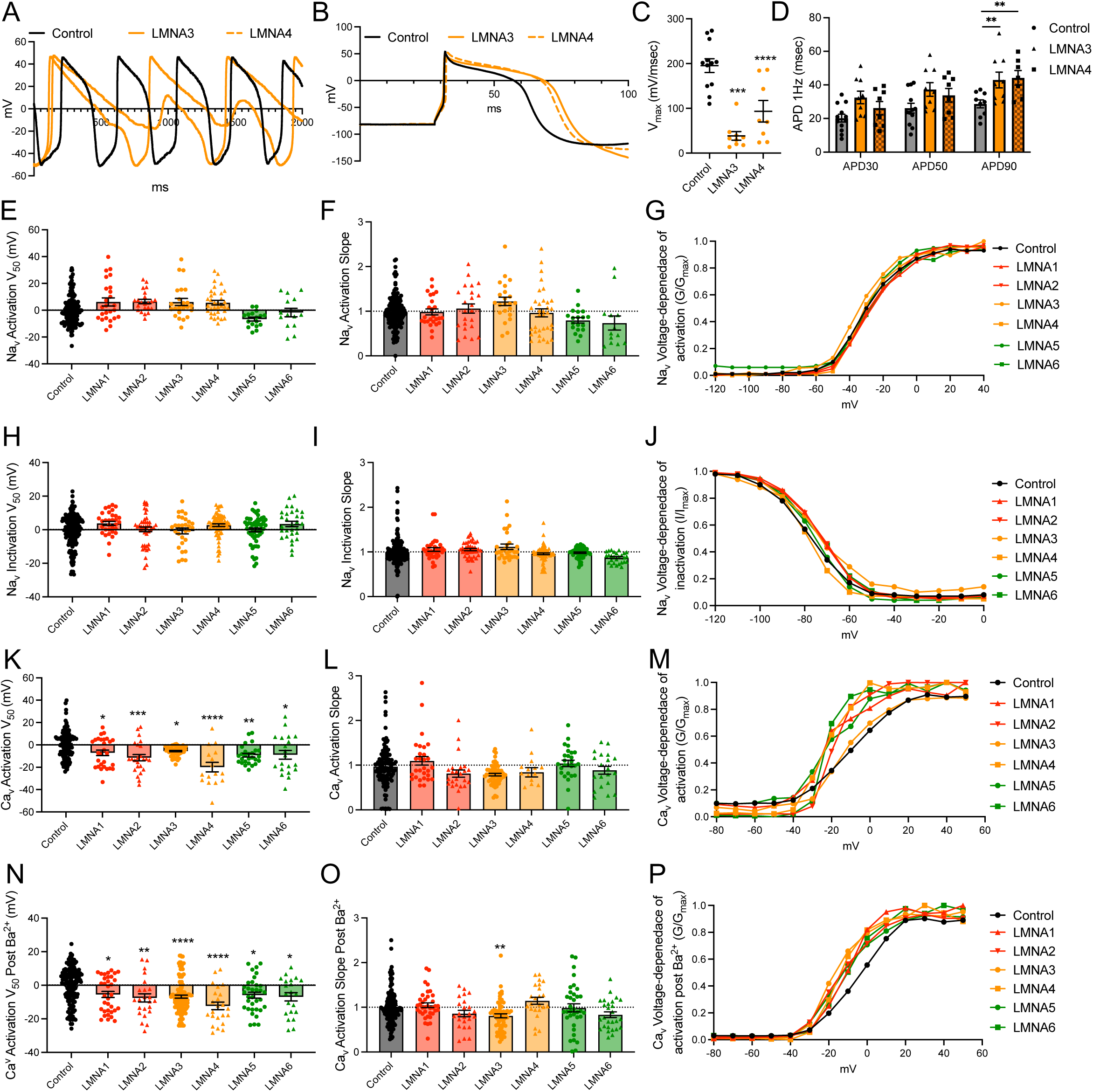
Manual and automated patch clamp analysis of *LMNA* hiPSC-CMs. Representative traces of spontaneous **(A)** and induced **(B)** action potential recordings of control, LMNA3 and LMNA4 hiPSC-CMs. Quantifications of **C)** V_max_ and **D)** action potential durations (Control N = 12, LMNA3 N = 9, LMNA4 N =8). Quantification of **E)** V_50_ and **F)** slope of Na^+^ channel activation measured using SyncroPatch (Control N = 142, LMNA1 N = 25, LMNA2 N = 22, LMNA3 N = 23, LMNA4 N = 34, LMNA5 N = 17, LMNA6 N = 14). **G)** Representative Na^+^ channel voltage dependance of activation curves. Quantification of **H)** V_50_ and **I)** slope of Na^+^ channel inactivation (Control N = 196, LMNA1 N = 32, LMNA2 N = 44, LMNA3 N = 29, LMNA4 N = 71, LMNA5 N = 55, LMNA6 N = 30). **J)** Representative Na^+^ channel voltage dependance of inactivation curves. Quantification of **K)** V_50_ and **L)** slope of Ca^2+^ channel activation (Control N = 119, LMNA1 N = 28, LMNA2 N = 24, LMNA3 N = 48, LMNA4 N = 16, LMNA5 N = 27, LMNA6 N = 20). **M)** Representative Ca^2+^channel voltage dependance of activation curves. Quantification of **N)** V_50_ and **O)** slope of Ca^2+^ channel activation with external Ba^2+^ (Control N = 138, LMNA1 N = 34, LMNA2 N = 23, LMNA3 N = 60, LMNA4 N = 23, LMNA5 N = 34, LMNA6 N = 21). **P)** Representative Ca^2+^channel voltage dependance of activation curves with external Ba^2+^. N represent the number of assessed cells. *, **, *** and **** indicate *P-*value < 0.05, 0.01, 0.001 and 0.0001 respectively.

Although insightful, the manual patch clamp system is very low throughput and labor intensive. To access a high throughput assay for testing Na^+^ and Ca^2+^ channels in hiPSC-CMs derived from laminopathy patients and controls, an automated patch clamp platform (SyncroPatch 768PE) was utilized. **Supplementary Figure 3A-C** shows the average current-voltage traces for Na^+^, Ca^2+^, and Ca^2+^ post Ba^2+^ treatment recordings. Analysis of the voltage-dependence of activation for Na^+^ channels revealed no significant differences between *LMNA* hiPSC-CMs and controls (**Fig. 3E-G**). Similarly, voltage-dependence of inactivation was not significantly different between laminopathy and control hiPSC-CMs (**Fig. 3H-J**). Unlike sodium currents, the calcium currents conductance-voltage relationships were significantly hyperpolarized for *LMNA* hiPSC-CMs, while the Ca^2+^ activation slope remained the same (**Fig. 3K-M**). When Ba^2+^ was used as the permeating ion to increase current amplitude, this observation was further replicated (**Fig. 3N-P)**. These data indicate an increased calcium flow in all the *LMNA* hiPSC-CMs either with Ca^2+^or Ba^2+^ as the main permeant ion, which is suggestive of a universal phenotype leading to arrhythmia in the patients.

### Calcium handling and the contractility are altered in laminopathy in laminopathy hiPSC-CMs

Since arrhythmia is one of the common complications of LMNA-DCM patients, and our gene expression analysis as well as current clamp recordings point to the involvement of calcium, two high throughput assays for analysis of calcium handling and contraction were established to identify the differences between cardiomyocyte function between patients and controls. Calcium transient recordings of *LMNA* hiPSC-CMs compared to controls revealed significant differences between all LMNA-DCM patients compared to controls (**Fig. 4A** and **Supplementary Fig. 4**). Further analysis of the calcium transient demonstrated significant prolongation of calcium transient duration 75% (CTD_75_), full width at half maximum (FWHM) and decay time from 75% to 25% (T_75-25_) in *LMNA* hiPSC-CMs compared to controls (**Fig. 4B**). This data is in line with our previous gene expression and current clamp analysis and identifies malfunction of calcium handling as a potential mechanism of arrhythmia in LMNA-DCM.

**Figure 4.**
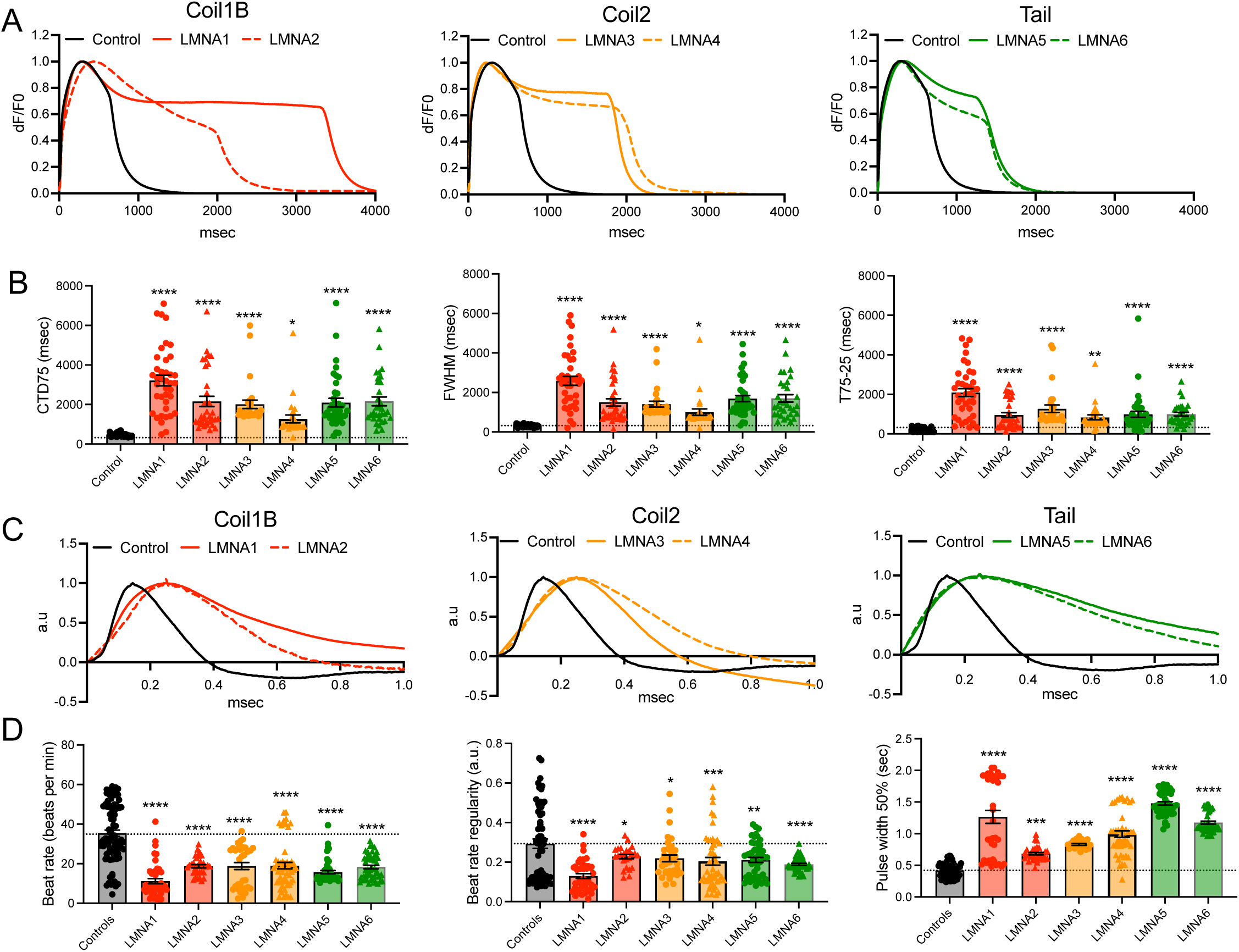
Calcium transient and contractility of *LMNA* hiPSC-CMs. **A)** Representative traces of calcium transients of *LMNA* hiPSC-CMs organised based on the location of their variants. **B)** Quantification of intracellular calcium traces (Control N = 53, LMNA1 N = 39, LMNA2 N = 35, LMNA3 N = 29, LMNA4 N = 26, LMNA5 N = 39, LMNA6 N = 32). **C)** Representative traces of impedance analysis of *LMNA* hiPSC-CMs organised based on the location of their variants. **D)** Quantification of impedance traces (Control N = 63, LMNA1 N = 37, LMNA2 N = 33, LMNA3 N = 20, LMNA4 N = 45, LMNA5 N = 50, LMNA6 N = 48). The dotted line indicates the average value for five control hiPSC-CMs. CTD_75_: calcium transient duration 75%, FWHM: full width at half maximum, T_75-25_: decay time from 75% to 25%. Data presented here are from at least three independent differentiations and the N number for calcium imaging indicates the number of assessed wells and for contractility analysis indicates the number of assessed sweeps. Data analysed using one way ANOVA. *, **, *** and **** indicate *P-*value < 0.05, 0.01, 0.001 and 0.0001 respectively.

In addition to calcium transients, we also studied the differences between *LMNA* and control hiPSC-CMs contractility inferred from impedance recordings. Similar to calcium transients, differences in the impedance traces were observed between the *LMNA* hiPSC-CMs and controls (**Fig. 4C**). Analysis of the beat rate suggested a significant decrease in the spontaneous beating velocity of all *LMNA* hiPSC-CMs compared to controls (**Fig. 4D**). Additionally, measurements of beat rate regularity indicated a significant decrease in *LMNA* hiPSC-CMs compared to controls (**Fig. 4D**). This finding is correlated with patients’ arrhythmia phenotype indicating it was successfully recapitulated in our *in vitro* model. Interestingly, pulse width at 50% was significantly greater in *LMNA* hiPSC-CMs (**Fig. 4D**). This observation was in line with our previous finding of the prolonged calcium transients in *LMNA* hiPSC-CMs. Additionally, it shows the applicability of our high throughput assays to successfully recapitulated LMNA-DCM phenotype *in vitro*.

### Correction of *LMNA* variants attenuates the abnormal *LMNA* hiPSC-CM phenotype *in vitro*

To determine whether the abnormal phenotypes observed in *LMNA* hiPSC-CMs were attributable to *LMNA* variants, representative mutations in the LMNA1, LMNA3, and LMNA5 lines were corrected using CRISPR/Cas9 genome editing. Corrected clones were established for each line, and Sanger sequencing confirmed precise correction of the respective *LMNA* variants (**Fig. 5A–C**). Functional analysis of hiPSC-CMs from the corrected lines, compared to their isogenic controls, demonstrated rescue of the abnormal calcium handling, as evidenced by a significant reduction in calcium transient duration parameters (CTD_75_, FWHM, and T_75–25_; **Supplementary Fig. 5A** and **Fig. 5D, 5G, 5J**). Impedance measurements further showed that *LMNA* correction resulted in a significant increase in beat rate and beat rate regularity, accompanied by a significant decrease in pulse width at 50% (**Supplementary Fig. 5B** and **Fig. 5E, 5H, 5K**). Morphological assessment of hiPSC-CMs from the corrected lines also revealed restoration of normal nuclear architecture, with a marked reduction in the number of cells exhibiting nuclear membrane blebbing (**Fig. 5F, 5I, 5L**). Collectively, these findings demonstrate that correction of *LMNA* variants restores both functional and morphological abnormalities in *LMNA* variant carrier lines, confirming that the *LMNA* mutations are causative of the observed phenotypes.

**Figure 5.**
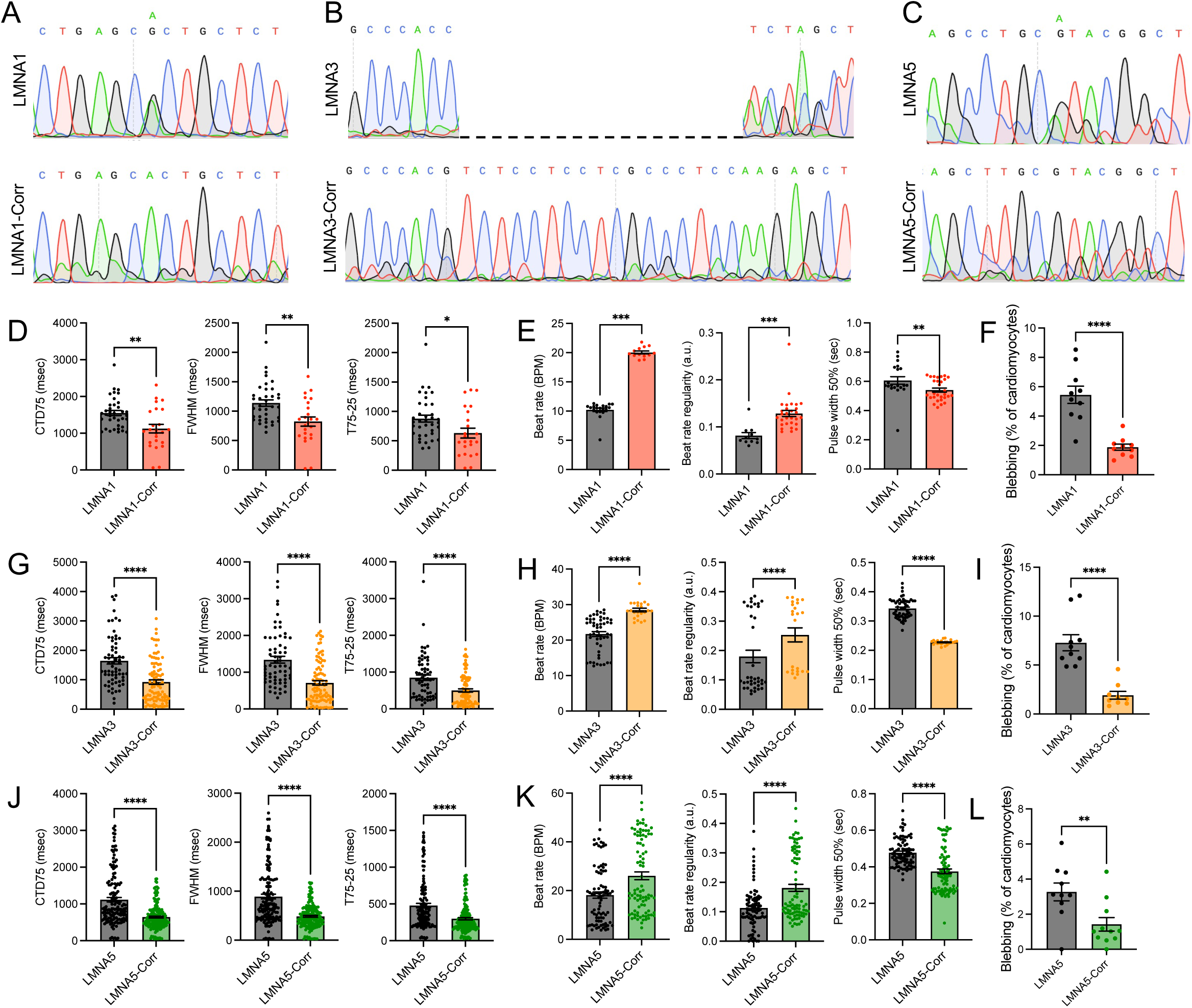
CRISPR/Cas9 correction of *LMNA* variants. Sanger sequencing of **A)** LMNA1, **B)** LMNA3, and **C)** LMNA5 before and after correction. **D)** Calcium transient (LMNA1 N = 37, LMNA1-Corr N = 24), **E)** impedance (LMNA1 N = 20, LMNA1-Corr N = 13), and **F)** nuclear membrane deformation (LMNA1 N = 10, LMNA1-Corr N = 10) analysis of LMNA1-Corrected hiPSC-CMs compared to LMNA1. **G)** Calcium transient (LMNA3 N = 64, LMNA3-Corr N = 94), **H)** impedance (LMNA3 N = 52, LMNA3-Corr N = 23), and **I)** nuclear membrane deformation (LMNA3 N = 10, LMNA3-Corr N = 9) analysis of LMNA3-Corrected hiPSC-CMs compared to LMNA3. **J)** Calcium transient (LMNA5 N = 150, LMNA5-Corr N = 195), **K)** impedance (LMNA5 N = 84, LMNA5-Corr N = 87), and **L)** nuclear membrane deformation (LMNA5 N = 10, LMNA5-Corr N = 11) analysis of LMNA5-Corrected hiPSC-CMs compared to LMNA5. CTD_75_: calcium transient duration 75%, FWHM: full width at half maximum, T_75-25_: decay time from 75% to 25%. Data presented here are from at least three independent differentiations and the N number for calcium imaging indicates the number of assessed wells, for contractility analysis indicates the number of assessed sweeps, and for blebbing the number of assessed fields of view. Data analysed using one way ANOVA. *, **, *** and **** indicate *P-*value < 0.05, 0.01, 0.001 and 0.0001, respectively.

### The abnormal calcium transients of *LMNA* hiPSC-CMs can be pharmacologically corrected

To assess whether the *LMNA* hiPSC-CMs respond to pharmacological treatments, cells were treated with sirolimus (rapamycin) which was previously reported to have positive outcomes in animal models of laminopathy^34,35^. hiPSC-CMs were treated with different dosages of sirolimus ranging from 10 nM to 100 μM and their calcium transients were studied. The 100 μM sirolimus was toxic for the assessed hiPSC-CMs as indicated by the absence of calcium signals. Our analysis demonstrated that for each assayed *LMNA* hiPSC-CMs, there was at least one concentration of sirolimus which could successfully correct all the abnormal calcium transient parameters (**Fig. 6** and **Supplementary Fig. 6**). Specifically, our analysis demonstrated that for LMNA1 treatment with 1 μM of sirolimus was able to correct all abnormal calcium transient parameters (**Fig. 6A**). For LMNA2, 10 nM, 100 nM and 1 µM concentrations were able to correct the increased CTD_75_ and FWHM and bring them down to the control level as indicated by nonsignificant differences in these parameters between treated LMNA2 hiPSC-CMs and control hiPSC-CMs. But only 1 µM of sirolimus corrected the T_75-25_ parameter (**Supplementary Fig. 6A).** For LMNA3 hiPSC-CMs, 10 nM of sirolimus was enough to correct all the abnormal calcium transient parameters, while 10 µM concentration only fixed the FWHM and T_75-25_ (**Fig. 6B**). Similarly, 1 and 10 µM of sirolimus corrected all the calcium parameters of LMNA4 (**Supplementary Fig. 6B).** For LMNA5, sirolimus corrected CTD_75_ at 10 nM, 100 nM and 10 µM; FWHM by 100 nM and 10 µM; and T_75-25_ by 10 µM (**Fig. 6C**). Finally, for LMNA6, all calcium parameters were corrected by 10 µM sirolimus treatment (**Supplementary Fig. 6C).** These data align with previous reports that suggested a positive effect of sirolimus in the treatment of mouse models of laminopathy^34,35^. However, the extensive side effects of sirolimus prohibits its application in clinic. Thus, it is crucial to search for alternative treatments.

**Figure 6.**
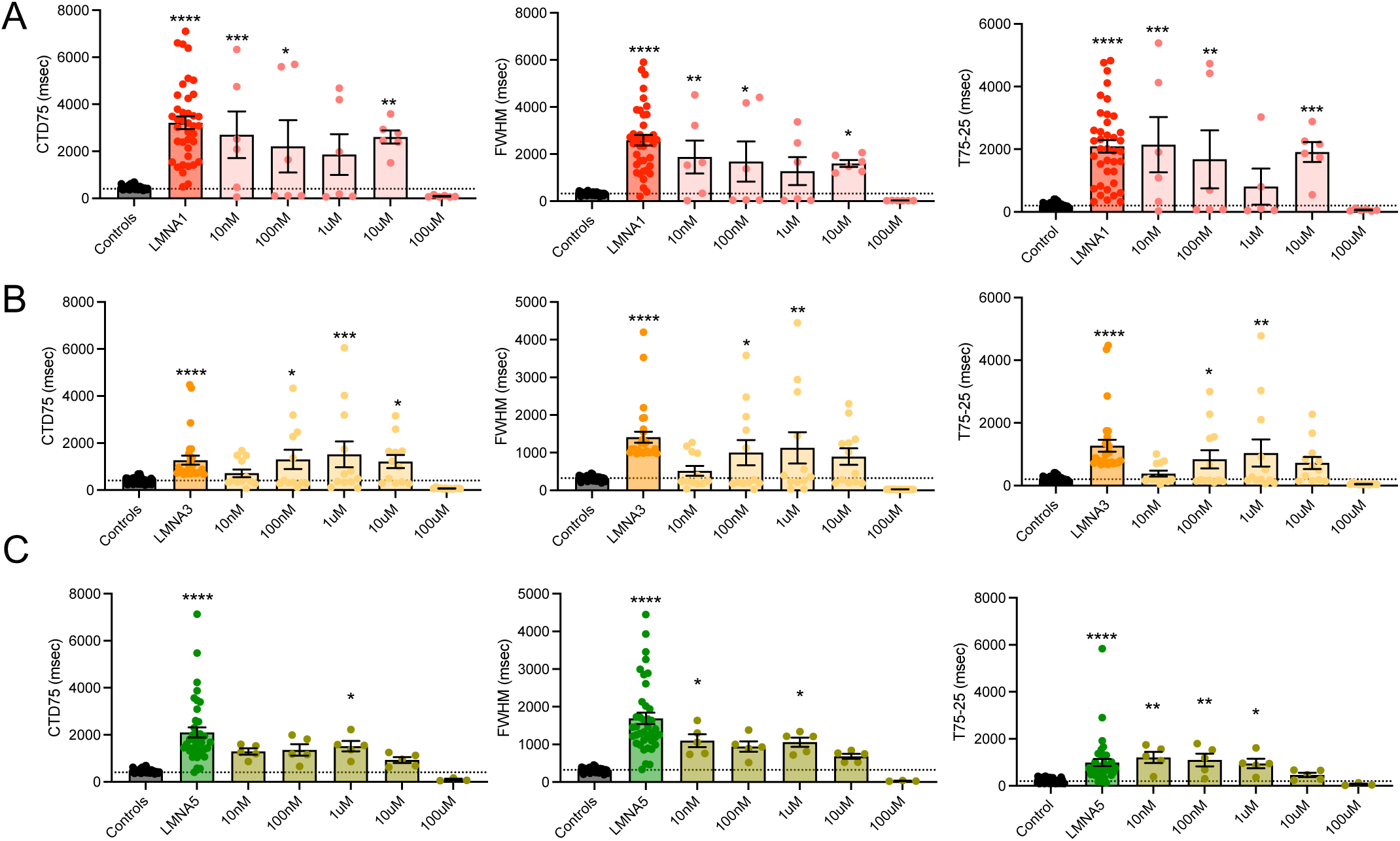
Calcium transient analysis of *LMNA* hiPSC-CMs post sirolimus treatment. Calcium transient analysis of **A)** LMNA1 (Control N = 53, LMNA1 N = 39, 10nM N = 6, 100nM N = 6, 1uM N = 6, 10uM N = 6, 100uM N = 6), **B)** LMNA3 (Control N = 53, LMNA3 N = 29, 10nM N = 12, 100nM N = 12, 1uM N = 12, 10uM N = 12, 100uM N = 12), and **C)** LMNA5 (Control N = 53, LMNA5 N = 39, 10nM N = 5, 100nM N = 5, 1uM N = 5, 10uM N = 5, 100uM N = 3) post treatment with different doses of sirolimus. The dotted line indicates the average value for five control hiPSC-CMs. CTD_75_: calcium transient duration 75%, FWHM: full width at half maximum, T_75-25_: decay time from 75% to 25%. Data presented here are from at least three independent differentiations and the N number for calcium imaging indicates the number of assessed wells and for contractility analysis indicates the number of assessed sweeps. Data analysed using one way ANOVA. *, **, *** and **** indicate *P-*value < 0.05, 0.01, 0.001 and 0.0001 respectively.

### Unbiased high-throughput drug screening reveals novel compounds for LMNA-DCM treatment

Our previous proof-of-concept experiment established the predictive validity of our *in vitro LMNA* hiPSC-CMs model for evaluating the impact of drug treatments. Next, to search for potential repurposed drug treatment for LMNA-DCM, we screened 1280 FDA approved drugs using Prestwick chemical library on all *LMNA* hiPSC-CMs. Drugs were screened based on their potential to successfully reduce all the prolonged calcium transient parameters including CTD_75_ **(Fig. 7A)**, FWHM (**Supplementary Fig. 7A**) and T_75-25_ (**Supplementary Fig. 7B**) to the control level. This functional screening revealed that for each tested *LMNA* hiPSC-CMs, there were specific drugs which were able to correct all the abnormal calcium transient parameters (**Fig. 7B**). Some of these compounds were unique to one *LMNA* hiPSC line while others were shared between two or more assessed *LMNA* cell lines (**Fig. 7C**). Further analysis revealed that out of 1280 drugs, 876 of them were effective on at least one *LMNA* hiPSC-CMs cell line (**Fig. 7D**). When considering drugs which were shared between two, three, four, five, or six *LMNA* hiPSC-CMs, this number decreased to 269, 41, 9, 2 and 1, respectively (**Fig. 7D**). The only compound that was shared between all the tested *LMNA* hiPSC-CMs was cyproheptadine hydrochloride.

**Figure 7.**
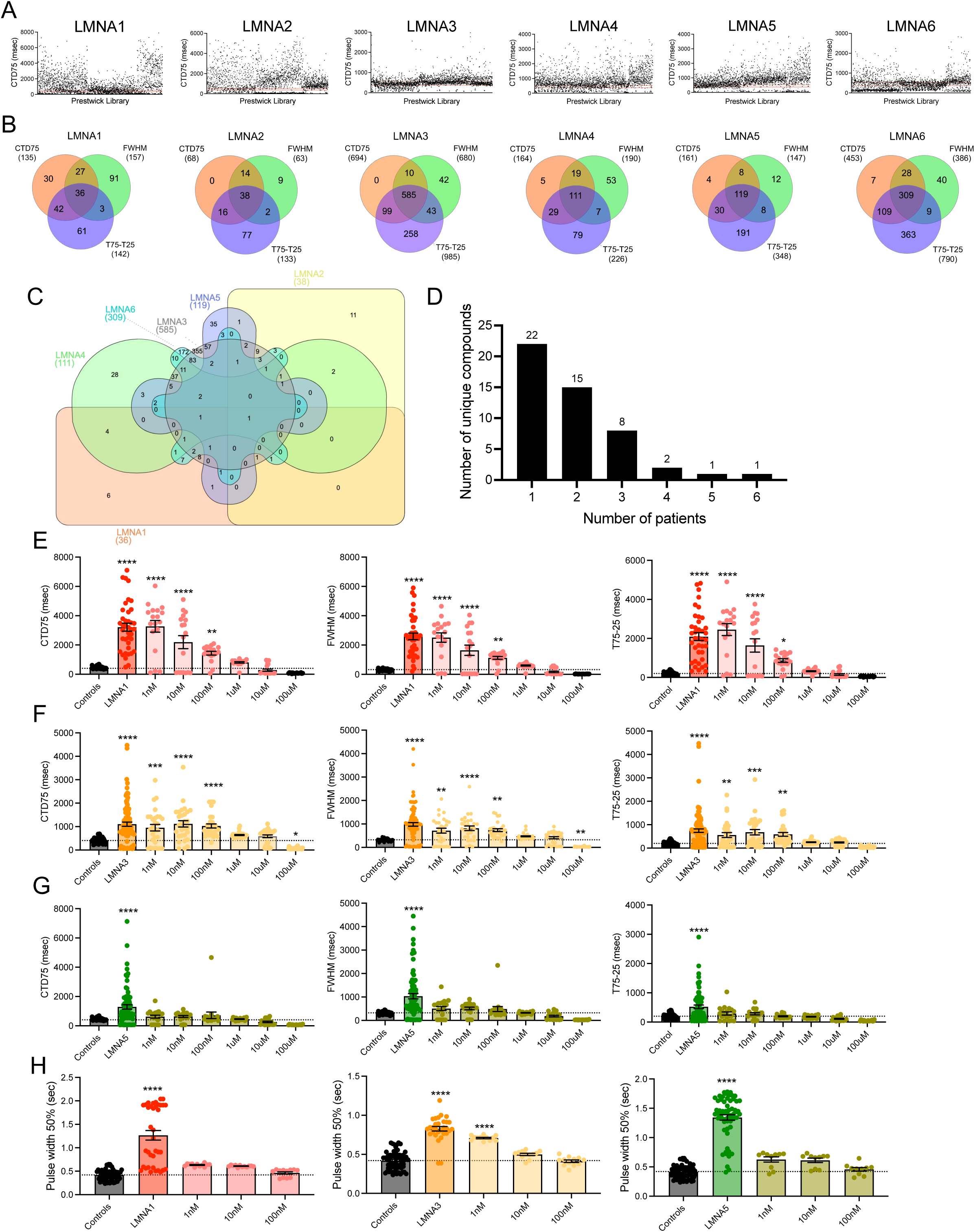
Unbiased high throughput drug screening and cyproheptadine response of *LMNA* hiPSC-CMs. **A)** Calcium transient duration 75% of *LMNA* hiPSC-CMs post treatment with Prestwich drug library. The two red dotted lines indicate the average of control CTD_75_ ± standard deviation. Each dot is the average of three different sets of experiments. **B)** Venn diagram presenting the overlap of the compounds from Prestwick drug library correcting calcium transient parameters of CTD_75_, FWHM and T_75-25_ for each *LMNA* hiPSC-CMs. **C)** Venn diagram presenting the overlap of the compounds alleviating impaired calcium transient phenotype for each *LMNA* hiPSC-CMs. **D)** The number of drugs that corrected all the impaired aspects of calcium transients that were shared between different number of laminopathy patients. Calcium transient analysis of **E)** LMNA1 (Control N = 53, LMNA1 N = 39, 1nM N = 18, 10nM N = 18, 100nM N = 18, 1uM N = 18, 10uM N = 18, 100uM N = 18), **F)** LMNA3 (Control N = 53, LMNA3 N = 95, 1nM N = 29, 10nM N = 29, 100nM N = 29, 1uM N = 29, 10uM N = 29, 100uM N = 29) and **G)** LMNA5 (Control N = 53, LMNA5 N = 77, 1nM N = 19, 10nM N = 19, 100nM N = 19, 1uM N = 19, 10uM N = 19, 100uM N = 19) post treatment with different doses of cyproheptadine. The N number indicates the number of assessed wells. **H)** Impedance analysis of LMNA1 (Control N = 63, LMNA1 N = 37, 1nM N = 12, 10nM N = 12, 100nM N = 12), LMNA3 (Control N = 63, LMNA3 N = 31, 1nM N = 12, 10nM N = 12, 100nM N = 12), and LMNA5 (Control N = 63, LMNA5 N = 60, 1nM N = 10, 10nM N = 10, 100nM N = 10) based on pulse width 50% post treatment with different doses of cyproheptadine. Data presented here are from at least three independent differentiations and the N number for calcium imaging indicates the number of assessed wells and for contractility analysis indicates the number of assessed sweeps. The dotted line indicates the average value for five control hiPSC-CMs. CTD_75_: calcium transient duration 75%, FWHM: full width at half maximum, T_75-25_: decay time from 75% to 25%. *, **, *** and **** indicate *P-*value < 0.05, 0.01, 0.001 and 0.0001 respectively.

### Cyproheptadine alleviates the impaired function of *LMNA* hiPSC-CMs

To further demonstrate the applicability of cyproheptadine treatment in mitigating the *LMNA* hiPSC-CM phenotype, *LMNA* hiPSC-CMs were exposed to different concentrations of cyproheptadine ranging from 1 nM to 100 mM for 72 h and then assessed based on their calcium transients and impedance. The highest concentration (100 mM) was toxic and was excluded from the analysis. This was further confirmed by measuring the cyproheptadine kill curve which demonstrated an LD_50_ of 16.15 ± 0.14 mM (**Supplementary Fig. 8A**). Our analysis revealed that for LMNA1, LMNA3 and LMNA6 the concentrations of 1 and 10 mM were effective to reduce the prolonged CTD_75_, FWHM and T_75-25_ to control level (**Fig. 7E-F** and **Supplementary Fig. 8D).** For LMNA4, 100 nM, 1 and 10 mM were effective (**Supplementary Fig. 8C).** Finally, for LMNA2 and LMNA5 treatment with cyproheptadine at all the measured concentrations (1 nM to 10 mM) reduced the abnormal calcium transients (**Fig. 7G** and **Supplementary Fig. 8B)**. To assess whether shortening the calcium transient is cyproheptadine general effect or it is a LMNA-DCM specific phenotype, a control hiPSC-CMs was treated with similar concentrations of cyproheptadine. Interestingly, no significant differences in the calcium transient parameters of the control hiPSC-CMs were observed post cyproheptadine treatment (**Supplementary Fig. 8E)**. This finding suggests that cyproheptadine does not affect the control hiPSC-CMs calcium transient.

Next, to test the effect of cyproheptadine treatment on other impaired functions of the LMNA-DCM, the contractile function of *LMNA* hiPSC-CMs was assessed post 72 h of treatment with different consecrations of the drug ranging from 1 nM to 100 nM. Like calcium transients, impedance analysis post cyproheptadine treatment demonstrated alleviation of the disease phenotype in all assayed *LMNA* hiPSC-CMs. Specifically, for LMNA 1, 4, 5 and 6 all 1, 10 and 100 nM were able to reduce the prolonged pulse width 50% down to control level (**Fig. 7H** and **Supplementary Fig. 9A)**. For LMNA2 100 nM and for LMNA3 and LMNA6 10 and 100 nM of cyproheptadine were enough to correct the abnormal pulse width 50% (**Fig. 7H** and **Supplementary Fig. 9A)**. Interestingly, similar to calcium transient measurements, treatment of healthy hiPSC-CMs with cyproheptadine did not change the contractility parameters (**Supplementary Fig. 9B**) which could strengthen a patient-specific response of cyproheptadine.

Finally, to harness the mechanism by which cyproheptadine rectifies the abnormal *LMNA* hiPSC-CMs phenotype, the expression of its direct targets in controls vs. *LMNA* hiPSC-CMs was studied (**Supplementary Fig. 10A**). Our analysis indicated that out of 18 known targets of cyproheptadine, 17 were very lowly expressed (< 10 TPM), thus were removed from further investigation. The only remaining target that was expressed in our hiPSC-CMs was *CHRM2* which was downregulated in *LMNA* hiPSC-CMs comparted to controls, albeit not reaching statistical significance. It is an interesting observation since the variants in *CHRM2* have previously been linked to DCM^36^. This might suggest that cyproheptadine alleviates the *LMNA* hiPSC-CMs functional phenotype through *CHRM2*. To assess this hypothesis, we sought to see whether there is an over representation of the drugs that target *CHRM2* directly from our functional drug screening. Our analysis revealed 46 approved drugs that target *CHRM2* directly. Out of this, only 27 were present in the Prestwick Chemical Library used for our drug screening. Out of these 27 drugs, 20 drugs (74%) had positive effect on at least one, 6 drugs (22%) on two or more, 2 drugs (7.4%) on two or more, and only one drug (3.7%), cyproheptadine, on four or more *LMNA* hiPSC-CMs (**Supplementary Fig. 10B**). This suggests that cyproheptadine might alleviates the abnormal function of laminopathy hiPSC-CMs through *CHRM2*.

## Discussion

The identification of reliable and effective therapies for *LMNA*-related dilated cardiomyopathy (LMNA-DCM) remains essential to improve patients’ quality of life. A human *in vitro* model that faithfully recapitulates patient-specific phenotypes offers the potential for personalized, functional drug testing. We demonstrate, for the first time, the applicability of patient-specific hiPSC-CMs in functional, high-throughput, unbiased drug screening. By utilizing a cohort of six patients carrying three distinct *LMNA* variants distributed across the gene, we were able to delineate both variant-specific and common disease features amenable to pharmacological targeting.

Our comprehensive characterization of *LMNA*-hiPSC-CMs uncovered several hallmark features: nuclear membrane deformation, altered electrophysiology, and impaired contractility, reflecting the arrhythmic phenotypes seen clinically. Gene expression profiling revealed downregulation of cardiac gene networks alongside activation of cardiomyopathy-associated pathways. Interestingly, *LMNA* itself was downregulated, while other intermediate filaments remained unchanged, suggesting a regulatory role of *LMNA* in maintaining the expression of its direct targets. Pathway enrichment analysis highlighted dysregulated calcium handling as a central abnormality, a finding supported by measurable prolongation of CTD_75_, FWHM, and T_75–25_ in calcium transient assays and accompanied by elevated pulse width 50% in impedance measurements across all patient lines. Current clamp analysis confirmed aberrant calcium channel function. Through CRISPR/Cas9-mediated correction of *LMNA* variants, we demonstrated that these functional and morphological abnormalities are causally linked to the mutations, as correction restored normal calcium handling, impedance parameters, and nuclear morphology. Collectively, these data point to intrinsic, cell-autonomous defects in cardiomyocyte physiology that likely drive arrhythmogenesis in LMNA-DCM, supporting the potential for variant-agnostic therapeutic targeting of calcium channel dysfunction.

Our findings align with and extend previous studies in *LMNA* and other cardiomyopathy models. Abnormal calcium handling has consistently been reported as an early and central event in disease progression, preceding overt ventricular dysfunction. For example, hiPSC-CMs carrying a troponin T mutation (R173W) showed increased calcium buffering, predisposing to arrhythmia^37^, while *Lmna*^H222P/H222P^ mice displayed elevated sarcolipin levels, inhibiting SERCA and contributing to abnormal Ca²⁺ transients^38^. More recently, hiPSC-CMs harboring *LMNA* frameshift mutations revealed reactive oxygen species (ROS)-driven degradation of SIRT1, leading to CaMKII and RyR2 activation and abnormal calcium release, a phenotype corrected by CRISPR correction^39^. In parallel, *LMNA* mutations have been shown to disrupt TRPV4-mediated calcium influx, with TRPV4 inhibition rescuing abnormal calcium loading^40^, and to activate MAPK and TGF-β pathways that exacerbate stress signaling and apoptosis^14^. Other studies highlight chromatin-level effects, showing that mutant *LMNA* perturbs lamin–chromatin interactions and causes misexpression of non-cardiac genes, suggesting that structural and transcriptional mechanisms may converge to produce the LMNA-DCM phenotype^41^. However, these studies have largely focused on single variants, limiting generalizability. By including three distinct variants across six patients, our study represents the largest LMNA-DCM hiPSC-CM cohort to date, allowing us to distinguish common versus variant-specific features and to demonstrate that calcium handling abnormalities are shared across genotypes.

To evaluate the translational potential of our platform, we performed a proof-of-concept pharmacological screen. Treatment with sirolimus, an mTOR inhibitor with previously reported benefits in DCM ^34,35^, improved calcium transients, validating the use of our hiPSC-CM system to identify candidate drugs. High-throughput functional assays revealed multiple compounds capable of rescuing calcium abnormalities, among which cyproheptadine emerged as the only drug effective across all variants. It is worth noting that the Prestwick Chemical Library® contains multiple compounds known to modulate calcium handling, including but not limited to milrinone, digoxin, dantrolene, verapamil, diltiazem, and nifedipine. However, none of these compounds reproduced the consistent correction observed with cyproheptadine, suggesting that its mechanism of action is not limited to a general enhancement of calcium cycling. This conclusion is further supported by our observation that cyproheptadine not only corrected the abnormal calcium-handling phenotype in *LMNA* hiPSC-CMs but also rescued contractility and reduced membrane blebbing in diseased cells. The variability in minimal effective concentrations across patient-specific LMNA hiPSC-CM lines likely reflects inter-individual differences in genetic background and disease severity, a known feature of hiPSC-based models, rather than nonspecific pharmacological activity, given the consistent functional rescue observed across all LMNA lines. Cyproheptadine, a first-generation H1 antihistamine with affinity for serotonin receptors and calcium channels, has been shown to reduce action potential duration in cardiac tissue^42^ and to prevent fibrotic remodeling after myocardial infarction^43^. In our study, physiological concentrations of cyproheptadine^44^ normalized CTD_75_, FWHM, T_75–25_, and pulse width 50% across all *LMNA*-hiPSC-CM lines. Interestingly, only *CHRM2*, a known cyproheptadine target, was highly expressed in our system, and *CHRM2* variants have been previously linked to cardiomyopathy^36^ , suggesting a plausible mechanism of action. Additionally, a recent study proposed ROS elevation as a potential mechanism by which *LMNA* variants contribute to DCM^39^. Notably, cyproheptadine has been suggested to mitigate this effect by scavenging ROS, thereby reducing oxidative stress and protecting cardiomyocytes^45^. These findings identify cyproheptadine as a potential therapeutic option for LMNA-DCM, with the advantage of broad efficacy independent of variant type. Future studies using cyproheptadine will elucidate the molecular and cellular mechanisms underlying its correction of the LMNA-associated DCM phenotype.

Patient-specific hiPSC-CMs provide a powerful human-based system for functional drug screening and precision therapy development. Their use bypasses interspecies differences inherent to animal models and enables direct testing of patient-relevant phenotypes. Nonetheless, current hiPSC-CMs remain immature compared to adult cardiomyocytes and lack the multicellular complexity of the human heart, which includes endothelial, fibroblast, and immune populations that contribute to disease progression^46^. Future directions should include multicellular cardiac organoids and engineered heart tissues, as well as integration of genome engineering platforms that enable systematic comparison of diverse *LMNA* variants within isogenic backgrounds^47^. Such approaches will provide deeper mechanistic insights and accelerate the development of targeted therapies for LMNA-DCM.

### Perspectives

#### Clinical Competency: Competency in Medical Knowledge

LMNA-associated dilated cardiomyopathy is characterized by high risk of arrhythmias and heart failure, yet its underlying mechanisms remain incompletely understood. This study identifies nuclear membrane abnormalities, reduced beat rate, arrhythmias, and prolonged calcium transients as shared disease mechanism across multiple *LMNA* variants, providing mechanistic insight that may inform future risk stratification and therapeutic approaches. Additionally, the use of patient-specific hiPSC-derived cardiomyocytes highlights the potential of human-based models to better predict disease phenotypes and drug responses compared to conventional systems.

#### Translational Outlook

Future studies are needed to validate these findings *in vivo* and in clinical settings to determine whether targeting calcium handling abnormalities can improve outcomes in *LMNA*-associated cardiomyopathy. The identification of cyproheptadine as a candidate therapeutic agent warrants further investigation in preclinical models and prospective clinical trials. More broadly, this work establishes a scalable platform for patient-specific drug discovery, which can be applied to other genetic cardiomyopathies to accelerate the development of precision therapies and improve translational success.

## Abbreviation

hiPSC: human induced pluripotent stem cells
hiPSC-CMs: human induced pluripotent stem cell-derived cardiomyocytes
DCM: dilated cardiomyopathy
LMNA-DCM: *LMNA*-related dilated cardiomyopathy
CTD_75_: calcium transient duration 75%
FWHM: full width at half maximum
T_75-25_: decay time from 75% to 25%
APD: action potential duration

